# Neurospora heterokaryons with complementary duplications and deficiencies in their constituent nuclei provide an approach to identify nucleus-limited genes

**DOI:** 10.1101/013482

**Authors:** Dev Ashish Giri, S. Rekha, Durgadas P. Kasbekar

## Abstract

Introgression is the transfer of genes or genomic regions from one species into another via hybridization and back-crosses. We have introgressed four translocations (*EB4*, *IBj5*, *UK14-1*, and *B362i*) from *Neurospora crassa* into *N. tetrasperma*. This enabled us to construct heterokaryotic [*T* + *N*] and [*Dp* + *Df*] strains in which the *mat-A* and *mat-a* nuclei have different genotypes. Self-crosses of the heterokaryons again produced [*T* + *N*] and [*Dp* + *Df*] progeny. From conidia (vegetative spores) produced by the heterokaryotic mycelia we obtained self-fertile (heterokaryotic) and self-sterile (homokaryotic) derivative strains. [*T* + *N*] heterokaryons produced homokaryotic derivatives of both mating types, but [*Dp* + *Df*] heterokaryons produced viable homokaryons of only the mating type of the *Dp* nucleus. All the four [*T* + *N*] heterokaryons, and three [*Dp* + *Df*] heterokaryons, produced both self-sterile and self-fertile conidial derivatives, but the [*Dp(B362i)* + *Df(B362i)*] heterokaryons produced only self-sterile ones. Conceivably, the *Df(B362i)* nuclei may be deleted for a nucleus-limited gene required for efficient mitosis or nuclear division, and whose deficit is not complemented by the neighboring *Dp(B362i)* nuclei. Repeat-induced point mutation (RIP) was shown to occur in a *Dp*-heterozygous cross, therefore RIP-alteration of a translocated segment would depend on relative numbers of self-crosses undergone in [*Dp* + *Df*] versus [*T* + *N*] ancestors.

## Introduction

Perkins (1997) described three elementary chromosome translocation (*T*) types in *Neurospora crassa*, namely, insertional (*IT*), quasiterminal (*QT*) and reciprocal (*RT*). *IT*s transfer a segment of a donor chromosome into a recipient chromosome without any reciprocal exchange, *QT*s transfer a distal segment of a donor chromosome to the tip of a recipient chromosome, distal to any essential gene, and presumably the donor chromosome breakpoint is capped with the tip from the recipient chromosome; and *RT*s reciprocally interchange the terminal segments of two chromosomes. Other chromosome rearrangements are essentially variants of these, e.g., an intrachromosomal transposition (*Tp*) is an *IT* in which the same chromosome is both donor and recipient, an inversion (*In*) is a *Tp* in which a chromosome segment is re-inserted in opposite orientation into the site from which it was derived, and there are complex rearrangements such as linked *RT* and *IT*. Three breakpoint junctions define an *IT*, viz, junction A created by the deletion on the donor chromosome, and junctions B and C (proximal and distal), created by the insertion into the recipient chromosome, whereas two breakpoint junctions define a *QT* or *RT*; junction A, between the breakpoint-proximal segment on the donor chromosome and the tip from the recipient chromosome, and junction B, between the breakpoint-proximal sequence on the recipient chromosome and the donor segment grafted onto it (Singh *et al*. 2010). In the cross of an *IT* or *QT* strain with normal sequence (*IT* x *N* or *QT* x *N*), alternate segregation produces eight parental-type ascospores (i.e., 4 *T* + 4 *N*), whereas adjacent 1 segregation generates non-parental ascospores, viz, four viable ascospores containing a duplication (*Dp*) of the translocated segment and four inviable ones with the complementary deficiency (*Df*). Viable ascospores blacken (B), whereas inviable ones remain white (W). Therefore, alternate and adjacent 1 segregation produce, respectively, 8B:0W and 4B:4W asci. Since both segregations are equally likely, *IT* x *N* and *QT* x *N* crosses are characterized by 8B:0W = 4B:4W, whereas isosequential crosses (i.e., *N* x *N* or *T* x *T*) produce mostly 8B:0W asci (Perkins 1997). Adjacent 1 segregation in *RT* x *N*, yields only inviable ascospores bearing complementary duplications and deficiencies (i.e., *Dp1*/*Df2* and *Dp2*/*Df1*), and the asci are 0B:8W. Therefore, 8B:0W = 0B:8W signals *RT* x *N*. *Dp* strains (the viable segregants from 4B:4W asci) are recognizable by the characteristic barren phenotype they impart to *Dp* x *N* crosses, wherein normal looking perithecia are made, but only a few exceptional ascospores are produced (Perkins 1997). Barrenness is caused by meiotic silencing by unpaired DNA (MSUD), an RNAi-mediated process that eliminates the transcripts of any gene not properly paired during meiosis with a homologue at an allelic position (Shiu *et al.* 2001). Presumably, *Dp*-borne genes, including those underlying ascus and ascospore development, fail to properly pair in a *Dp* x *N* cross, and their silencing by MSUD renders the cross barren. The breakpoint junctions of several *IT*s, *QT*s, and *RT*s were defined in our laboratory (Singh 2010; Singh *et al*. 2010). PCR with breakpoint junction-specific primers can now be used to distinguish the *Dp* progeny from their *T* and *N* siblings. *IT* progeny contain all three breakpoints (A, B, and C), *Dp* progeny contain B and C, but not A, and *N* progeny contain none. While *Dp*s have been extensively studied (Perkins 1997; Kasbekar 2013), the use of *Df*s was limited to flagging the *Dp*-bearing 4B:4W asci. We now report the generation of [*Dp* + *Df*] heterokaryons with complementing duplications and deficiencies in their constituent nuclei. They were obtained by introgressing *N. crassa IT*s (and a *QT*) into *N. tetrasperma*. Introgression is the transfer of genes or genomic regions from one species into another (Rieger *et al.* 1991).

Eight ascospores form per ascus in *N. crassa*, whereas four are formed in *N. tetrasperma*. In both species the parental *mat A* and *mat a* nuclei fuse in the ascogenous cell to produce a diploid zygote nucleus that immediately undergoes meiosis and a post-meiotic mitosis to generate eight haploid nuclei (4 *mat A* + 4 *mat a*). In *N. crassa* the nuclei are partitioned into the eight initially uninucleate ascospores (Raju 1980). In contrast, *N. tetrasperma* ascospores are initially binucleate, receiving a pair of non-sister nuclei (1 *mat A* + 1 *mat a*) (Raju and Perkins 1994). Thus, *N. crassa* ascospores produce homokaryotic mycelia that are either *mat A* or *mat a* in mating type, whereas *N. tetrasperma* ascospores can produce heterokaryotic mycelia with nuclei of both mating types. In *N. crassa*, a sexual cross perforce requires mycelia from two different ascospores, one *mat A*, the other *mat a*, thus making the lifecycle “heterothallic”; whereas a heterokaryotic *N. tetrasperma* mycelium from a single ascospore bearing nuclei of both mating types is competent to undergo a self-cross, making the lifecycle “pseudohomothallic”. However, a subset of conidia (vegetative spores) produced by a heterokaryotic *N. tetrasperma* mycelium can be homokaryotic by chance, and *N. tetrasperma* ascogenesis occasionally produces five or more (upto eight) ascospores, by replacement of one or more dikaryotic ascospore by a pair of smaller homokaryotic ones (Raju 1992). The dominant *Eight-spore* (*E*) mutant increases the frequency of such replacement, although *E*-homozygous crosses are infertile (Calhoun and Howe 1968). Mycelia from homokaryotic conidia or ascospores can cross with like mycelia of the opposite mating type. Therefore, *N. tetrasperma* is actually a facultatively heterothallic species.

Here, we have introgressed three *N. crassa IT*s (*EB4*, *IBj5*, and *B362i*) and one *QT* (*UK14-1*) into *N. tetrasperma*, and shown that *T* x *N* crosses produce both [*T* + *N*] and [*Dp* + *Df*] heterokaryotic progeny. We found that unlike the other heterokaryons, the [*Dp(B362i)* + *Df(B362i)*] heterokaryons produced only self-sterile conidia, possibly because the *Df(B362i)* nuclei are missing a putative “nucleus-limited” gene required for efficient mitosis and nuclear division. Additionally, we show that a *Dp*-heterozygous cross can exhibit RIP (repeat-induced point mutation), the sexual stage-specific process that induces G:C to A:T mutations in duplicated DNA (Selker 1990).

### Materials and Methods

#### Neurospora strains and general genetic manipulations

All Neurospora strains were obtained from the Fungal Genetics Stock Center, University of Missouri, Kansas City, Missouri, USA, unless otherwise indicated. *N. crassa*: OR *A* (FGSC 987) and OR *a* (FGSC 988), are the standard laboratory Oak Ridge strains; the translocation strains *T*(*VR ->VII*)*EB4 A* (FGSC 3046), *T(VIL->IR) IBj5 cpc-1 A* (FGSC 4433), *T(VIR > VL) UK14-1 A* (FGSC 6958), and *T(IV->I)B362i A* (FGSC 2935), (abbreviated to *T(EB4) A*, *T(IBj5)*, *T(UK14-1)*, and *T(B362i)*). *T(EB4)*, *T(IBj5)*, and *T(B362i)* are *IT*s, whereas *T(UK14-1)* is a *QT*. These translocations have been described by Perkins (1997) and Singh (2010).

The semi-dominant MSUD suppressor strains *Sad-1 A* (FGSC 8740) and *Sad-1 a* (FGSC 8741), were gifted by Robert L. Metzenberg and are described by Shiu *et al*. (2001). The *sad-1* locus encodes an RNA-dependent RNA polymerase essential for MSUD, and the *Sad-1* suppressor allele is presumed to prevent proper pairing of its wild-type homologue thus inducing it to autogenously silence itself (Shiu *et al*. 2001). The MSUD testers *pan-2; his-3::his-3^+^ Bml^r^ A* (FGSC 8755); *pan-2; his-3::his-3^+^ Bml^r^ a* (FGSC 8756); *pan-2; his-3::his-3^+^ mei-3^+^ A* (FGSC 8759); and *pan-2; his-3::his-3^+^ mei-3^+^ a* (FGSC 8760) (hereafter designated as *:: Bml^r^ A*,:: *Bml^r^ a*, *::mei-3 A*, and *::mei-3 a*) are described by Raju *et al*. (2007). Another tester, *rid his-3; VIIL::r^ef2^-hph A* (ISU3117) (hereafter *::r^ef2^*) was a gift from Dr. Tom Hammond (University of Missouri). The *::Bml^r^* and *::mei-3* testers have an extra copy of the *bml* (β-tubulin) or *mei-3* gene inserted ectopically in the *his-3* locus in chromosome 1, while the tester strain *::r^ef2^* has a copy of the *r* (*round spores*) gene inserted ectopically into chromosome 7. In a cross of the tester with wild type, the ectopic copy remains unpaired in meiosis and results in elimination of all its homologous transcripts, including from the paired endogenous copies. In crosses of *::Bml^r^*, *::mei-3* and *::r^ef2^* with the wild type, the *bml*, *mei-3*, and *r* genes, respectively, are silenced. Silencing of the *bml* or *mei-3* gene arrests normal ascus development (Raju *et al*. 2007; Kasbekar *et al*. 2011), and silencing of *r* causes all eight ascospores to be round instead of the normal spindle shaped. Homozygous *tester A* x *tester a* crosses do not show MSUD, nor do crosses of the testers with the *Sad-1* suppressor of MSUD, and the asci developed normally (Raju *et al*. 2007; Kasbekar *et al*. 2011).

*N. tetrasperma*: the standard strains 85 *A* (FGSC 1270) and 85 *a* (FGSC 1271); the *E* mutants *lwn; al(102), E A* (FGSC 2783) and *lwn; al(102), E a* (FGSC 2784) (hereafter *E A* and *E a*). *N. crassa* / *N. tetrasperma* hybrid strain: *C4,T4 a* (FGSC 1778). The *C4,T4 a* strain has four *N. crassa* great-grandparents and four *N. tetrasperma* great-grandparents (Metzenberg and Ahlgren 1969). The *N. crassa* great-grandparents were of the OR background, whereas the *N. tetrasperma* great-grandparents were of the 343.6 *A E* background (Metzenberg and Ahlgren 1969).

Neurospora genetic analysis was done essentially as described by Davis and De Serres (1970). Metzenberg’s (2003) alternative recipe was used for making Medium N.

#### Outline of the introgression crosses and characterization of the resultant strains

Crosses between *N. crassa* and *N. tetrasperma* strains are almost completely sterile. However, both *N. crassa* strain OR *A* and *N. tetrasperma* strain 85 *A* can cross with the *N. crassa* / *N. tetrasperma* hybrid strain *C4,T4 a* and produce viable progeny (Perkins 1991; also see Table 3 of this paper) therefore we used the *C4,T4 a* strain as a bridging strain for the initial introgression crosses. The *N. crassa T* strains were crossed with *C4T4 a* and *T* progeny from these crosses (designated *T^1xC4T4^*) were distinguished from their *Dp* and *N* siblings by PCR with breakpoint junction-specific primers. Nominally, 50% of the genome of *T^1xC4T4^* progeny is derived from the *C4,T4 a* parent. The *T^1xC4T4^A* strains were crossed with *C4,T4 a*, to obtain the *T^2xC4T4^* progeny in a like manner. Crosses of *T^2xC4T4^* with the opposite mating type derivative of strain *85* were productive, and their *T* progeny were designated as *T^1x85^*. Likewise, *T^1x85^* x *85* yielded *T^2x85^*, etc. After two to three iterations of the crosses with *85*, we recovered progeny ascospores that produced mycelium of dual mating specificity characteristic of *N. tetrasperma*. That is, the resulting mycelium could cross with both 85*A* and *a*, and it could also undergo a self-cross. A heterokaryotic strain containing all three breakpoints (A, B and C) is potentially of genotype [*T* + *N*] or [*Dp* + *Df*].

The [*T* + *N*] and [*Dp* + *Df*] heterokaryons are distinguishable, since the former produces homokaryotic conidial derivatives of both mating types, whereas the latter produces viable homokaryons of only the mating type of the *Dp* nucleus. Conidia from self-fertile heterokaryotic strains were streaked onto Vogel’s–FGS medium, and well-isolated conidial germlings were transferred to SCM to distinguish self-fertile (heterokaryotic) from self-sterile (homokaryotic) conidial derivatives. The mating type of the self-sterile conidial derivatives was determined by crossing to the single mating type derivatives *85 a* and *85 A*. If all the self-sterile conidial derivatives are of a single mating type, then the heterokaryon from which they were derived is likely [*Dp* + *Df*], else it is [*T* + *N*]. The results were confirmed by PCR with primers for the breakpoint junctions and *mat* ideomorphs and DNA of the homokaryotic conidial derivatives.

Four homokaryotic *T* type conidial derivatives from the self-fertile [*T* + *N*] heterokaryons, namely, *T(EB4)^Nt^ a* from the heterokaryon 3E1 (Table 2 serial number 3), *T(IBj5)^Nt^ a* from I4 (Table 2 serial number 18), *T(UK14-1)^Nt^ a* from U9 (Table 2 serial number 24), and *T(B362i)^Nt^A* from 19B7 (Table 2 serial number 35) were used in the experiments whose results are summarized in Tables 3 and 4.

### Results

#### *N. tetrasperma* [*T* + *N*] and [*Dp* + *Df*] strains can switch genotype via self-crosses

The salient features of the *N. crassa* translocations *T(EB4)*, *T(IBj5)*, *T(UK14-1)*, and *T(B362i)* are summarized in Table 1, along with accession numbers of their breakpoint junction sequences. The introgression of these translocations into *N. tetrasperma* is outlined in the “Materials and Methods” section, and Figure 1 schematically presents the actual crosses done.

**Figure 1.**
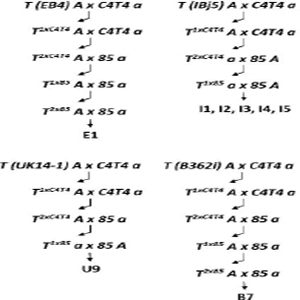
Introgression crosses. *T(EB4) A*, *T(IBj5) A*, *T(UK14-1) A*, and *T(B362i) A* strains of *N. crassa* were crossed with the *C4T4 a* hybrid strain. Bent arrows represent PCR with breakpoint junction-specific primers to distinguish the translocation progeny (e.g., *T^1xC4T4^*) from their *Dp* and *N* siblings. *T^1xC4T4^A* x *C4T4 a* yielded *T^2xC4T4^ A* or *T^2xC4T4^ a* strains, which were productive in crosses with opposite mating type homokaryotic derivatives of *N. tetrasperma* strain *85*. *T^1x85^* progeny were crossed with *85 a* or *85 A* to obtain the self-fertile heterokaryotic strains I1-I5 (for *IBj5*) and U9 (for *UK14-1*), or the *T^2x85^* strains (for *EB4* and *B362i*). Crosses of *T^2x85^* with *85 a* or *85 A* produced the heterokaryons E1 and B7. From self-cross of the heterokaryons we obtained self-fertile progeny that were genotyped as [*T* + *N*] or [*Dp* + *Df*] (see Table 2).

**Table 1.** Translocations used in this study.

| Translocation | Size<br>(bp) | Genes<br>(N) | Breakpoint Junction Sequence<br>(Accession Number) |  |  |
| --- | --- | --- | --- | --- | --- |
|  |  |  | A | B | C |
| *T(VR > VII) EB4* | 145282 | 39 | GQ504681 | GQ504682 | GQ504683 |
| *T(VIL > IR) IBj5* | 405319 | 120 | GQ504684 | GD504685 | - |
| *T(VIR > VL) UK14-1* | 490958 | 126 | GQ504703 | - | NA |
| *T(IVR > I) B362i* | 118782 | 36 | GQ504697 | GQ504698 | GQ504699 |

NA- not applicable.

Introgression of *T(EB4)* yielded the self-fertile heterokaryotic strain designated E1 (Figure 1). Using E1 genomic DNA as template and *T(EB4)* breakpoint junction-specific primers, all three breakpoint junctions (A B and C) of *T(EB4)* could be amplified by PCR (data not shown). A heterokaryon possessing all the three breakpoints is potentially of genotype [*T(EB4)* + *N*] or [*Dp(EB4)* + *Df(EB4)*]. Heterokaryons of genotype [*T(EB4)* + *T(EB4)*], [*T(EB4)* + *Dp(EB4)*], or [*T(EB4)* + *Df(EB4)*] also fulfill this criterion, but, they were deemed to be less likely since one or more crossover is required to generate them. Eight self-fertile progeny from the self-cross of E1 were analyzed and the results, summarized in Table 2 (serial numbers 1-8), established that progeny 2E1, 4E1 and 6E1 were of the [*T(EB4)* + *N*] genotype, whereas 1E1, 9E1 and 12E1 were [*Dp(EB4)* + *Df(EB4)*]. Only six self-sterile conidial derivatives were obtained for 3E1, and PCR revealed all to be *Ta* type, suggesting that 3E1 has the genotype [*T a* + *Df A*]. However, given the small numbers tested, and possibility of skewed segregation, our results do not exclude the [*T a* + *N A*] genotype. No self-sterile conidial derivatives were obtained from 13E1, therefore its genotype was not determined.

**Table 2.**
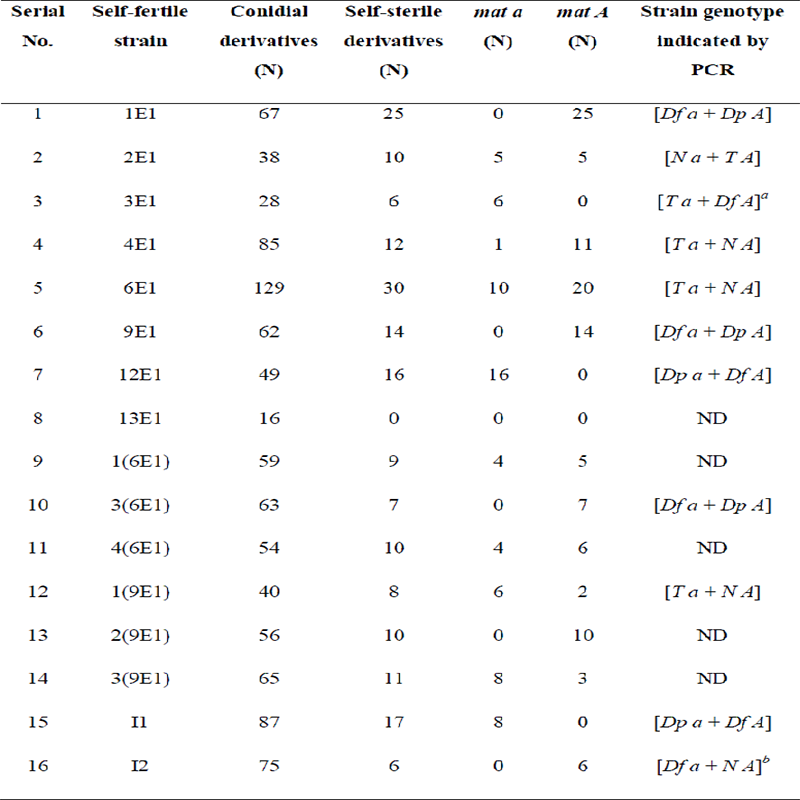
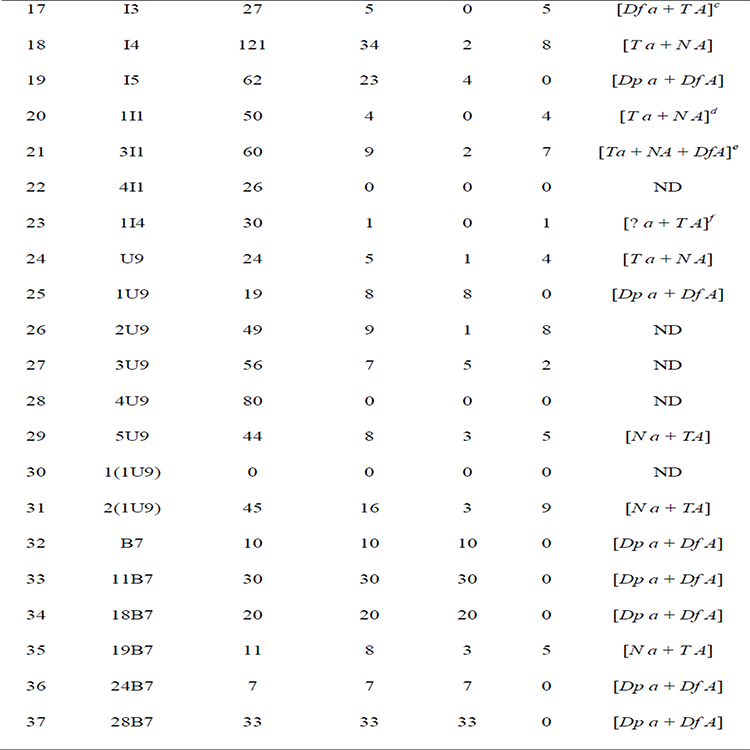
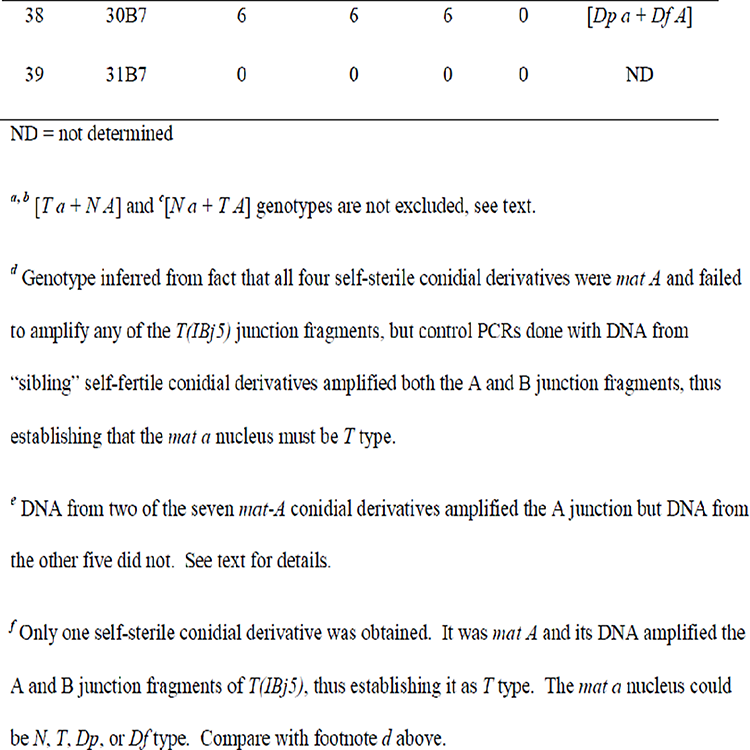
Genotype of self-fertile strains.

| **Serial**<br>**No.** | **Self-fertile**<br>**strain** | **Conidial**<br>**derivatives**<br>**(N)** | **Self-sterile**<br>**derivatives**<br>**(N)** | ***mat a***<br>**(N)** | ***mat A***<br>**(N)** | **Strain genotype**<br>**indicated by**<br>**PCR** |
| --- | --- | --- | --- | --- | --- | --- |
| 1 | 1E1 | 67 | 25 | 0 | 25 | [*Df a + Dp A*] |
| 2 | 2E1 | 38 | 10 | 5 | 5 | [*N a + T A*] |
| 3 | 3E1 | 28 | 6 | 6 | 0 | [*T a + Df A*]<sup>a</sup> |
| 4 | 4E1 | 85 | 12 | 1 | 11 | [*T a + N A*] |
| 5 | 6E1 | 129 | 30 | 10 | 20 | [*T a + N A*] |
| 6 | 9E1 | 62 | 14 | 0 | 14 | [*Df a + Dp A*] |
| 7 | 12E1 | 49 | 16 | 16 | 0 | [*Dp a + Df A*] |
| 8 | 13E1 | 16 | 0 | 0 | 0 | ND |
| 9 | 1(6E1) | 59 | 9 | 4 | 5 | ND |
| 10 | 3(6E1) | 63 | 7 | 0 | 7 | [*Df a + Dp A*] |
| 11 | 4(6E1) | 54 | 10 | 4 | 6 | ND |
| 12 | 1(9E1) | 40 | 8 | 6 | 2 | [*T a + N A*] |
| 13 | 2(9E1) | 56 | 10 | 0 | 10 | ND |
| 14 | 3(9E1) | 65 | 11 | 8 | 3 | ND |
| 15 | I1 | 87 | 17 | 8 | 0 | [*Dp a + Df A*] |
| 16 | I2 | 75 | 6 | 0 | 6 | [*Df a + N A*]<sup>b</sup> |
| 17 | I3 | 27 | 5 | 0 | 5 | $[Df\ a + TA]^c$ |
| 18 | I4 | 121 | 34 | 2 | 8 | $[Ta + NA]$ |
| 19 | I5 | 62 | 23 | 4 | 0 | $[Dp\ a + DfA]$ |
| 20 | 1I1 | 50 | 4 | 0 | 4 | $[Ta + NA]^d$ |
| 21 | 3I1 | 60 | 9 | 2 | 7 | $[Ta + NA + DfA]^e$ |
| 22 | 4I1 | 26 | 0 | 0 | 0 | ND |
| 23 | 1I4 | 30 | 1 | 0 | 1 | $[? \ a + TA]^f$ |
| 24 | U9 | 24 | 5 | 1 | 4 | $[Ta + NA]$ |
| 25 | 1U9 | 19 | 8 | 8 | 0 | $[Dp\ a + DfA]$ |
| 26 | 2U9 | 49 | 9 | 1 | 8 | ND |
| 27 | 3U9 | 56 | 7 | 5 | 2 | ND |
| 28 | 4U9 | 80 | 0 | 0 | 0 | ND |
| 29 | 5U9 | 44 | 8 | 3 | 5 | $[Na + TA]$ |
| 30 | 1(1U9) | 0 | 0 | 0 | 0 | ND |
| 31 | 2(1U9) | 45 | 16 | 3 | 9 | $[Na + TA]$ |
| 32 | B7 | 10 | 10 | 10 | 0 | $[Dp\ a + DfA]$ |
| 33 | 11B7 | 30 | 30 | 30 | 0 | $[Dp\ a + DfA]$ |
| 34 | 18B7 | 20 | 20 | 20 | 0 | $[Dp\ a + DfA]$ |
| 35 | 19B7 | 11 | 8 | 3 | 5 | $[Na + TA]$ |
| 36 | 24B7 | 7 | 7 | 7 | 0 | $[Dp\ a + DfA]$ |
| 37 | 28B7 | 33 | 33 | 33 | 0 | $[Dp\ a + DfA]$ |
| 38 | 30B7 | 6 | 6 | 6 | 0 | $[Dp\ a + Df\ A]$ |
| 39 | 31B7 | 0 | 0 | 0 | 0 | ND |

ND = not determined
<sup>a, b</sup> $[T\ a + N\ A]$ and <sup>c</sup> $[N\ a + T\ A]$ genotypes are not excluded, see text.
<sup>d</sup> Genotype inferred from fact that all four self-sterile conidial derivatives were *mat A* and failed to amplify any of the *T(IBj5)* junction fragments, but control PCRs done with DNA from “sibling” self-fertile conidial derivatives amplified both the A and B junction fragments, thus establishing that the *mat a* nucleus must be *T* type.
<sup>e</sup> DNA from two of the seven *mat-A* conidial derivatives amplified the A junction but DNA from the other five did not. See text for details.
<sup>f</sup> Only one self-sterile conidial derivative was obtained. It was *mat A* and its DNA amplified the A and B junction fragments of *T(IBj5)*, thus establishing it as *T* type. The *mat a* nucleus could be *N*, *T*, *Dp*, or *Df* type. Compare with footnote *d* above.

From self-crosses of strains 6E1 and 9E1 (see above) we examined 39 and 24 progeny, and found, respectively, 12 and nine were self-fertile. From a subset of self-fertile progeny we obtained self-sterile (i.e., homokaryotic) conidial derivatives and determined their mating type by crossing with 85 *a* and 85 *A*. Of three self-fertile progeny tested from strain 6E1, two appeared to be [*T(EB4) a* + *N A*] and one was [*Dp(EB4) a* + *Df(EB4) A*]; and of three self-fertile progeny tested from strain 9E1, one appeared to be [*Dp(EB4) a* + *Df(EB4) A*] and the other two to be [*T(EB4) a* + *N A*]. We used PCR to confirm the genotype of the [*Dp(EB4) a* + *Df(EB4) A*] progeny of 6E1, and of one [*T a* + *N A*] progeny of 9E1 (Table 2, serial numbers 10 and 12). These results showed that self-crosses of both [*T(EB4)* + *N*] and [*Dp(EB4)* + *Df(EB4)*] heterokaryons can again generate [*T(EB4)* + *N*] and [*Dp(EB4)* + *Df(EB4)*] progeny.

The translocations *T(IBj5)*, *T(UK14-1)*, and *T(B362i)* were introgressed in a like manner (Figure 1), and we recovered heterokaryons of genotype [*T(IBj5)* + *N*] and [*Dp(IBj5)* + *Df(IBj5)*] (Table 2, serial numbers 15, 18, 19, 20); [*T(UK14-1)* + *N*] and [*Dp(UK14-1)* + *Df(UK14-1)*] (Table 2, serial number 24, 25, 29, and 31); and [*T(B362i)* + *N*] and [*Dp(B362i)* + *Df(B362i)*] (Table 2, serial numbers 32-38). Self-crosses of the [*T* + *N*] and [*Dp* + *Df*] heterokaryons again produced progeny of the alternative genotype. Specifically, the [*Dp(IBj5)* + *Df(IBj5)*] type strain I1, produced the [*T(IBj5)* + *N*] type progeny strain 1I1 (Table 2, serial numbers 15 and 20); the [*T(UK14-1)* + *N*] type strain U9, produced the [*Dp(UK14-1)* + *Df(UK14-1)*] type progeny strain 1U9, whose self-cross, in turn, produced the [*T(UK14-1)* + *N*] type strain 2(1U9) (Table 2, serial numbers 24, 25, and 31); and the [*Dp(B362i)* + *Df(B362i)*] type strain B7, produced the [*T(B362i5)* + *N*] type progeny strain 19B7 (Table 2, compare serial numbers 32 and 35). In sum, our results show that [*T* + *N*] and [*Dp* + *Df*] genotypes are interchangeable through self-crosses.

The genotype of two heterokaryons was found to be putatively [*Df(IBj5) a* + *N A*] and [*Df(IBj5) a* + *T(IBj5)A*] (Table 2, serial numbers 16 and 17), but again we cannot rule out the possibility that skewed segregation in small numbers might account for the absence of self-sterile *T* or *N* conidial types, respectively, from what in fact might be [*T* + *N*] heterokaryons. One heterokaryon was found to contain three nuclear types, and its genotype was [*T(IBj5) a* + *NA* + *Df(IBj5)A*] (Table 2, serial number 21). It got flagged because two of the seven *mat A* self-sterile conidial derivatives examined possessed only the A, but not the B, junction of *T(IBj5)*, whereas the other five did not possess either junction. A homokaryon with only the A junction is not expected to be viable because it contains the *Df* chromosome, therefore we presume the two self-sterile derivatives were [*NA* + *Df(IBj5)A*] heterokaryons, therefore we infer that the genotype of the self-fertile strain was [*T(IBj5) a* + *NA* + *Df(IBj5)A*]. We defer to the discussion section a consideration of how such a strain might have arisen.

#### [*Dp(B362i)* + *Df(B362i)*] heterokaryons yield only self-sterile conidial derivatives

We expected the search for self-sterile homokaryotic conidial derivatives (above) to inevitably identify self-fertile ones as well. Indeed, for both [*T* + *N*] and [*Dp* + *Df*] strains of *EB4*, *IBj5*, and *UK14-1*, a majority of the conidial derivatives were self-fertile, and self-fertile conidial derivatives were also obtained from [*T(B362i)* + *N*] (Table 2, serial number 35). Therefore, we were very surprised to find that all 113 conidial derivatives from the six [*Dp(B362i)* + *Df(B362i)*] heterokaryons examined were self-sterile (Table 2, serial numbers 32-34 and 36-38).

As a control, we performed the cross *Dp(B362i) a* x *85 A* and 39 of 61 progeny tested were self-fertile. We examined ~10 conidial derivatives from each of 10 self-fertile progeny, and in every case at least three were self-fertile, and among the self-steriles we found both mating types. This suggested that unlike [*Dp(B362i)* + *Df(B362i)*], the [*Dp(B362i)* + *N*] heterokaryons make self-fertile conidia. A consideration of the implications of the [*Dp(B362i)* + *Df(B362i)*] heterokaryon’s phenotype is deferred to the discussion section.

#### Characterizing the *T* type homokaryons

The homokaryotic *T* type conidial derivatives designated as *T(EB4)^Nt^*, *T(IBj5)^Nt^*, *T(UK14-1)^Nt^*, and *T(B362i)^Nt^* were obtained from the self-fertile [*T* + *N*] heterokaryons (see Materials and Methods) and found to behave like bona fide *N. tetrasperma* strains. That is, their crosses with opposite mating type derivatives of *N. tetrasperma* strain *85* were fertile, whereas their crosses with *N. crassa* OR strains of opposite mating type were as infertile as the interspecies OR x *85* cross (Table 3). Control crosses of the *N. crassa T* strains (*T^Nc^*) with the OR strains of the opposite mating type were productive, but the crosses of the *T^Nc^* strains with the opposite mating type strain *85* derivatives were sterile (Table 3). The *C4,T4 a* hybrid strain produced viable ascospores in crosses with both OR *A* and *85 A* (Table 3).

**Table 3.** Ascospore productivity in crosses of *T* strains with OR and 85.

| Strain* | Productivity in cross |  |
| --- | --- | --- |
|  | x OR | x 85 |
| *T (EB4)<sup>Nc</sup>* | ++++ | - |
| *T (IBj5)<sup>Nc</sup>* | ++++ | - |
| *T (UK14-1)<sup>Nc</sup>* | ++++ | - |
| *T (B362i)<sup>Nc</sup>* | ++++ | - |
| *T (EB4)<sup>Nt</sup>* | - | ++++ |
| *T (IBj5)<sup>Nt</sup>* | - | ++++ |
| *T (UK14-1)<sup>Nt</sup>* | - | ++++ |
| *T (B362i)<sup>Nt</sup>* | - | ++++ |
| OR A | ++++ | - |
| 85 A | - | ++++ |
| *C4,T4 a* | +++ | ++ |

Ascospore yield: - < 100; 100 < ++ < 500; 500 < +++ < 5000; +++++ > 5000.
\* *T<sup>Nc</sup>* are *N. crassa T* strains, and *T<sup>Nt</sup>* the translocation breakpoint-bearing homokaryons derived from self-fertile heterokaryons.

To obtain larger numbers of eight-spored asci, we crossed the *T* strains with *E* strains of the opposite mating type. The *T(B362i)^Nt^* x *E* crosses were infertile, and *T(IBj5)^Nt^* x *E* produced mostly white inviable ascospores. However, *T(EB4)^Nt^* x *E* and *T(UK14-1)^Nt^* x *E* were productive, and the 8:0 and 4:4 ascus types were produced at comparable frequencies (Table 4), which is characteristic of *IT* x *N* and *QT* x *N* crosses in *N. crassa*.

**Table 4.** Ascus (octets) types from *T* x *N* crosses in *N. crassa* (*Nc*) and *N. tetrasperma* (*Nt*).

| Cross | Octets<br>(N) | Ascus type (%) |  |  |  |  |
| --- | --- | --- | --- | --- | --- | --- |
|  |  | 8:0 | 6:2 | 4:4 | 2:6 | 0:8 |
| *T<sup>Nc</sup> x OR* |  |  |  |  |  |  |
| *T (EB4)<sup>Nc</sup>* | 56 | 56 | 5 | 36 | 3 | 0 |
| *T (IBj5)<sup>Nc</sup>* | 72 | 31 | 43 | 19 | 6 | 1 |
| *T (UK14-1)<sup>Nc</sup>* | 86 | 21 | 46 | 24 | 7 | 4 |
| *T (B362i)<sup>Nc</sup>* | 83 | 55 | 13 | 32 | 0 | 0 |
| *T<sup>Nt</sup> x E* |  |  |  |  |  |  |
| *T (EB4)<sup>Nt</sup>* | 96 | 32 | 29 | 40 | 2 | 0 |
| *T (UK14-1)<sup>Nt</sup>* | 101 | 47 | 15 | 37 | 1 | 1 |

#### RIP in a *Dp*-heterozygous *N. tetrasperma* cross

In *N. crassa*, crosses involving *Dp* strains can generate RIP-induced mutant progeny (Perkins *et al*. 1997). Therefore, we expected crosses of *N. tetrasperma Dp* strains also would yield RIP-induced mutants. *Dp(EB4)* duplicates the *ad-7* (*adenine requiring-7*) gene (Perkins 1997). Ascospores from *Dp(EB4) a* x *E A* were germinated on adenine-supplemented Vogel’s-FGS medium, 130 germlings were picked to adenine-supplemented Vogel’s-glucose medium, and then their growth was tested on unsupplemented Vogel’s-glucose medium. Three adenine-requiring auxotrophic strains were identified among 125 progeny examined. One was a heterokaryon, but the other two were *N* type homokaryons. In both the homokaryons, the *ad-7* locus was altered by several RIP mutations (G:C to A:T transitions) (Table 5).

**Table 5.** RIP-induced *ad-7* mutants from *Dp(EB4)* x *E*.

|  | *Nt* | *Nc* | *RIP1*<sup>2</sup> | *RIP2*<sup>3</sup> |
| --- | --- | --- | --- | --- |
| **85 A**<sup>1</sup> | 6 (4) | 69 (27) | 45 (39) | 40 (34) |

Comparison of *ad-7* gene sequences from 85 A (*N. tetrasperma* FGSC 1270 <sup>1</sup>KP006652); *Nt* (*N. tetrasperma* FGSC 2508 sequence ID GL891302, nucleotides 1022253 to 1024228); *Nc* (*N. crassa* OR74A FungiDB database (fungidb.org) NCU04216, nucleotides 5238499 to 5240480); and the RIP-induced mutants *RIP1* (<sup>2</sup>KP006653) and *RIP2* (<sup>3</sup>KP006654). Superscripts indicate accession numbers of sequences determined in this work. Numbers in parenthesis indicate nucleotide differences in the exons, and altered amino acid residues are listed below in bold. The mutants exhibit several nonsense and missense mutations.
85 A /*Nt* (4) - T40T, S216S, I306I, F475F
85 A /*Nc* (27) - T40T, F94F, S132S, D156D, D189D, R201R, S216S, L221L, F229F, L233L, G245G, Y264Y, I306I, D377D, A396A, A404A, A421A, P423P, V445V, A456A, P462P, F475F, G506G, T522T, **E524D**, I538I, H541H
85 A / *RIP1* (39) - **H22Y**, N97N, **H106Y**, I111I, **L120F**, **Q122\***, T130T, D131D, N139N, **R222H**, **H249Y**, **Q250\***, A268A, **P270T**, T272T, I273I, S391S, A396A, R399R, I402I, A404A, P409P, I410I, I411I, A421A, **P423L**, **Q424\***, L426L, **H429Y**, D435D, H439H, I440I, **Q448\***, **H464Y**, **Q465\***, T507T, N508N, V518V, **H541Y**
85 A / *RIP2* (34) - **Q49\***, Y72Y, N97N, G101G, I111I, **L120F**, **H127Y**, V153V, S203S, Y212Y, **Q241\***, **Q245\***, **Q250\***, **Q254\***, A291A, **L298F**, V303V, I309I, T315T, **S324L**, V341V, F345F, I346I, A363A, G383G, T384T, A396A, R399R, **H414Y**, N534N, **H541Y**, N542N, S546S

#### MSUD is relatively weak in crosses involving *C4,T4* a and *85 A/a*

Ramakrishnan *et al*. (2011) classified wild-isolated *N. crassa* strains into three types based on the strength of MSUD in their crosses with tester strains, namely, “OR” (which includes the standard laboratory strains OR *A* and OR *a*), “Sad” (which includes the semi-dominant *Sad* suppressors of MSUD obtained from the OR background), and “Esm”. MSUD was strongest in crosses with the OR type, of intermediate strength in crosses with the Esm type, and weakest in crosses with the Sad type. We performed crosses of *C4,T4 a* with the MSUD tester strains. The results summarized in Supplementary Figure 1 show that *C4,T4 a* partially suppresses MSUD like the Esm type strains. Presumably, the Esm phenotype of *C4,T4 a* is derived from its 343.6 *A E* ancestor.

In *N. crassa*, crosses of *Dp(EB4)* and *Dp(IBj5)*strains with the OR type strains were barren, whereas their crosses with the Sad type strains were fertile (Ramakrishnan *et al*. 2011). Most Esm type strains gave a fertile cross with *Dp(EB4)* and a barren cross with *Dp(IBj5)*, although some Esm type gave barren crosses with both the *Dp*s, and fewer still gave fertile crosses with them. Other results of Ramakrishnan *et al*. (2011) suggested that *N. tetrasperma* strain *85* is Esm type. We crossed the *N. tetrasperma Dp(EB4)* and *Dp(IBj5)* strains with opposite mating type derivatives of strain *85* and found the crosses were as productive as those of *T(EB4)* and *T(IBj5)* with strain *85* (data not shown). These results are consistent with the classification of strain *85* as Esm or Sad type.

### Discussion

We have constructed self-fertile heterokaryotic [*T* + *N*] and [*Dp* + *Df*] Neurospora strains by introgressing *N. crassa* translocations into *N. tetrasperma*. Both types of heterokaryons when self-crossed again produced [*T* + *N*] and [*Dp* + *Df*] progeny. The [*T* + *N*] and [*Dp* + *Df*] mycelia were distinguishable because the former yielded self-sterile homokaryotic derivatives of both mating types, while the latter produced homokaryons of only the mating type of the *Dp* component. Our success in constructing the heterokaryons depended on the unambiguous identification of the translocation progeny (*T*) from each round of introgression crosses by PCR with breakpoint junction-specific primers, as represented by the bent arrows in Figure 1. In 1984, N. B. Raju crossed a *N. crassa* rearrangement strain with the giant-spore *Banana* (*Ban*) mutant with the hope that he could rescue the [*Dp* + *Df*] heterokaryon. He had previously used this approach to rescue *Sk^S^* nuclei from a *Sk^S^* x *Sk^K^* cross in [*Sk^S^* + *Sk^K^*] heterokaryotic ascospores (Raju 1979), but analysis of progeny nuclei in the mixed cultures from the later experiment was not easy, and his efforts were inconclusive (personal communication from Dr. N. B. Raju to DPK). D. D. Perkins also alluded to his obtaining [*Dp* + *Df*] heterokaryons (Perkins 1997), but his results remained unpublished. Therefore, to the best of our knowledge, this is the first report of heterokaryons with complementary duplications and deficiencies in their constituent nuclei.

The introgressions were facilitated by use of the *C4,T4 a* bridging strain (Perkins 1991). We found that *C4,T4 a* is a moderate suppressor of MSUD (Supplementary Figure 1). Shiu *et al*. (2001) had shown that the *N. crassa* MSUD suppressor *Sad-1* can partially breach the *N. crassa* / *N. tetrasperma* interspecies barrier. Although *C4,T4 a* does not suppress MSUD as strongly as *Sad-1 a*, its moderate suppressor phenotype might contribute to increase ascospore production in crosses with both *N. crassa* and *N. tetrasperma* strains. The *N. crassa T* strains were first crossed with *C4T4 a*, and *T A* progeny from this cross were crossed with *C4T4 a*. *T* progeny from the latter crosses were used to initiate two or three additional rounds of crosses with opposite mating type derivatives of *N. tetrasperma* strain *85*. Although this sequence of introgression crosses can be continued indefinitely, we stopped once they began yielding self-fertile progeny. Nominally, one nucleus in the B7, E1, I1, I2, I3, I4, I5, and U9 self-fertile progeny of Figure 1 derives 1/8 or 1/4 of its genome from the non-*85* genetic background (*N. crassa T* or *C4T4 a*), and, depending on the translocation, this fraction is enriched for sequences from the translocation chromosome (1, 4, 5, 6, or 7). Therefore, in principle, by sequencing one can identify strain *85* genome segments that are replaceable by non-*85* without losing the pseudohomothallic life-cycle.

Self-crosses of the [*T* + *N*] / [*Dp* + *Df*] strains also appeared to produce [*T* + *Df*] and [*N* + *Df*] genotypes. These genotypes can be produced by a breakpoint-proximal crossover on the donor or recipient chromosome followed by second-division segregation of the resulting *Df*- or *N*-bearing chromosome into both *mat a* and *mat A* nuclei. Breakpoint-proximal crossover produces 6:2 ascus types in *IT* x *N* and *QT* x *N* crosses in *N. crassa* (Perkins 1997), and in *IT* x *E* and *QT* x *E* crosses in *N. tetrasperma* (Table 4). Second-division segregation can also generate [*Ta* + *TA*], [*Dp a* + *Dp A*], [*N a* + *N A*], [*N* + *Dp*], and [*T* + *Dp*] progeny types. One self-fertile heterokaryon had the genotype [*T a* + *NA* + *Df A*]. This triple heterokaryon could have arisen from an interstitial cross-over, that in *N. crassa* would have produced a 6:2 tetratype ascus with two nuclei each of the *T a*, *N A*, *Dp a*, and *Df A* genotypes (see Figure 1 in Perkins 1997). Presumably, three nuclei (*T a*, *N A*, and *Df A*) instead of two were partitioned during ascus development into the ascospore from which this strain was derived. In *N. crassa* crosses heterozygous for the *Fsp-1* (*Four-spore-1*) or *Fsp-2* (*Four-spore-2*) mutant, rare two- and three-spored asci are occasionally formed that contain heterokaryotic ascospores (Raju 1986; Perkins *et al*. 2001). Presumably, our [*T a* + *NA* + *Df A*] strain had a similar provenance.

A *Dp*-borne gene (*ad-7*) was shown to be alterable by RIP. Since RIP occurs in self-cross of [*Dp* + *Df*] but not [*T* + *N*], RIP-alteration of a translocated segment would depend on the number of adjacent 1 versus alternate segregations in its ancestral crosses.

All the four [*T* + *N*] heterokaryons, and three of the [*Dp* + *Df*] heterokaryons, yielded both self-fertile and self-sterile conidial derivatives. The proportion of self-fertile derivatives from the [*T(B362i)* + *N*] tested was < 50%, but for the other six heterokaryons it was > 50%. However, the [*Dp(B362i)* + *Df(B362i)*] heterokaryons appeared to be an exception, in that, none of the six strains tested produced any self-fertile conidial derivatives. One hypothesis to account for these results is that *Df(B362i)* nuclei divide much less efficiently than *Dp(B362i)* nuclei, resulting in a rapid dwindling of their number in the [*Dp(B362i)* + *Df(B362i)*] heterokaryon. Consequently, they are less likely to be packaged along with the *Dp(B362i)* nuclei during conidiation, when typically three to ten (presumably randomly picked) nuclei are partitioned into each conidium. Even if a few heterokaryotic [*Dp(B362i)* + *Df(B362i)*] conidia form, the *Df(B362i)* nuclei are less likely to divide upon conidial germination, thus biasing even the heterokaryotic conidia to produce homokaryotic *Dp(B362i)* germlings. This hypothesis provides a plausible (and exonerating) explanation for an otherwise perturbing set of results that we had obtained in the early stages of this study. A strain was initially scored as [*Dp(B362i)* + *Df(B362i)*], because of its self-fertility and the ability of its DNA to support PCR amplification of all three *B362i*-specific junction fragments, but later it behaved like a *Dp(B362i)* homokaryon, because it was now self-sterile and its newly isolated DNA failed to amplify the A fragment (Dev Ashish Giri, unpublished results). A self-cross would selectively restore the *Df(B362i)* nuclear fraction back to 50%, but self-crosses are not possible once the *Df(B362i)* nuclei are completely lost.

In our model, the putative *Df(B362i)-*nuclear division defect occurs in a heterokaryotic cytoplasm, hence the gene presumed to be required for efficient division must have a null-phenotype that is non-complementable by the wild-type allele in the neighboring *Dp(B362i)* nuclei. A gene whose null allele (Δ) is not complemented by the wild-type allele (*WT*) in a [Δ + *WT*] heterokaryon can be considered to be nucleus-limited in function (Kasbekar, 2014). Although no nucleus-limited genes have been reported as yet, their existence in fungi is not ruled out, especially given the putative nucleus-limited behavior of the *N. crassa scon^c^* mutant (Burton and Metzenberg 1972), the DNA damage checkpoint signal in *Saccharomyces cerevisiae* (Demeter *et al*. 2000), and the MatIS gene silencing process in *Aspergillus nidulans* (Czaja *et al.* 2013). The *S. cerevisiae* precedent is germane to our model because it demonstrates that of two nuclei sharing the same cytoplasm, the one with damaged DNA arrests in mitosis without impeding progression through mitosis of the other with undamaged DNA.

Introgression of additional *N. crassa* translocations into *N. tetrasperma* holds out the prospect of uncovering more genotypes with putative nucleus-limited effects. Alternatively, *N. tetrasperma* can now be transformed (Kasbekar 2015), therefore a quicker way to screen for nucleus-limited genes might be via engineering of targeted integration of yeast *Frt* sites into the different chromosome arms, and using FLP recombinase to induce crossover and produce defined *RT*s. The [*Dp1*/*Df2* + *Dp2*/*Df1*] and [*RT* + *N*] heterokaryons generated in *RT* x *N* would enable us to screen two (or more) *Df*s in a single experiment.

## Acknowledgements

B. Navitha provided technical assistance. DAG was supported by a CSIR-UGC Junior Research Fellowship. DPK holds the Haldane Chair of the Centre for DNA Fingerprinting and Diagnostics (CDFD). This work was supported by a grant from the Department of Science and Technology, Government of India, and by CDFD Core Funds to DPK.

---

**Supplementary Figure 1.** *C4,T4 a* is a weak MSUD suppressor. Ascus development in crosses of the *N. crassa* strains OR *a* and *Sad-1 a*, and the *N. crassa* / *N. tetrasperma* hybrid strain *C4,T4 a* with the MSUD tester strains *::bml A*, *::mei-3 A* and *::r^ef2^*. Meiotic silencing of the *bml* (β-tubulin) and *mei-3* genes in the crosses with OR *a* disrupts ascus development, whereas its suppression in the crosses with *Sad-1 a* allows normal ascus development. Silencing of *r* in the cross with OR *a* causes all eight ascospores to be round, and its suppression by *Sad-1 a* restores the normal spindle shape. Silencing is evident in crosses of *C4,T4 a* with *::bml A* and *::r^ef2^A* but not in the cross with *::mei-3 A*. Partial suppression of MSUD by *C4,T4 a* is characteristic of Esm type strains (Ramakrishnan *et al*., 2011).

